# Rapid event-related, BOLD, NHP: choose two out of three

**DOI:** 10.1101/2020.01.27.919555

**Authors:** Vassilis Pelekanos, Robert M. Mok, Olivier Joly, Matthew Ainsworth, Diana Kyriazis, Maria G. Kelly, Andrew H. Bell, Nikolaus Kriegeskorte

## Abstract

Human functional magnetic resonance imaging (fMRI) typically employs the blood-oxygen-level-dependent (BOLD) contrast mechanism. In non-human primates (NHP), contrast enhancement is possible using monocrystalline iron-oxide nanoparticles (MION) contrast agent, which has a more temporally extended response function. However, using BOLD fMRI in NHP is desirable for interspecies comparison, and the faster response of the BOLD signal promises to be beneficial to rapid event-related (rER) designs. Here, we used rER BOLD fMRI in macaque monkeys while viewing real-world images, and found visual responses and category-selectivity consistent with previous studies. However, activity estimates were very noisy, suggesting that the lower contrast-to-noise ratio of BOLD, suboptimal behavioural performance, and motion artefacts, in combination, render rER BOLD fMRI challenging in NHP. Previous studies have shown that rER monkey fMRI is possible with MION, despite its prolonged response function. To understand this, we conducted simulations of the BOLD and MION response during rER designs, and found that no matter how fast the design, the greater amplitude of the MION response outweighs the contrast loss caused by greater temporal smoothing. We conclude that although any two of the three elements (rER, BOLD, NHP) have been shown to work well, the combination of all three is particularly challenging.

## Introduction

Functional magnetic resonance imaging (fMRI) has enabled the acquisition of whole-brain images of brain activity in humans and other animals. The technique has been used to functionally localize brain regions, with particular success in localizing regions selective for different visual categories, including face-, body-, object-, and place-selective areas in humans^1-3^ and non-human primates (NHP)^4-8^.

Human fMRI studies typically use the endogenous contrast agent deoxyhemoglobin, and measure the blood-oxygen-level-dependent (BOLD) signal. BOLD has been used in humans with a wide variety of experimental designs, including rapid event-related designs that give researchers great flexibility. In particular, rapid event-related fMRI enables condition-rich designs intended for pattern-information analyses^9^. BOLD fMRI has also been utilised by NHP studies (see below), however, many NHP studies have used the exogenous contrast agent monocrystalline iron oxide nanoparticle (MION) to increase the sensitivity of the measured signal. MION reflects blood volume, rather than blood oxygenation. Vanduffel et al.^10^ compared the use of BOLD vs. MION in block-design experiments in awake macaque monkeys to map the brain areas selective for motion. Their results not only matched monkey electrophysiology and human fMRI results, but also showed greater spatial localization and contrast increase in MION relative to BOLD. More recently, block-designs combined with MION have been predominantly used to localize fMRI-defined category-selective areas in macaques (for example^4,7,8,11^).

MION’s slower haemodynamic response is unproblematic in the context of block designs. Leite et al.^12^ compared MION with BOLD in macaques using visual checkerboard stimuli with varying presentation durations. They found that MION increased the functional sensitivity for stimuli presented at long durations, but brief or rapidly repeated stimulus presentations led to a greater attenuation of the signal compared to BOLD, consistent with a linear model capturing the dispersion of the response over time. This suggests that MION might be less sensitive for rapid event-related designs, whose high-temporal-frequency effects might not pass through the low-temporal-frequency filter of the MION response. However, event-related designs have been successfully used in MION fMRI studies previously^13,14^.

To understand the functional homologies and analogies between the human and the NHP brain, it would be desirable to use the same contrast mechanism in both species. Given that administering MION is an invasive procedure, not approved for routine use in humans, BOLD in NHPs might be the best approach for interspecies comparisons. Indeed, Pinsk et al.^5,6^ investigated visual category-selectivity using BOLD with block-designs, whereas BOLD in an event-related design with long delays was used by Kagan et al.^15^ and more recently by Kaskan et al.^16^. Interestingly, the faster temporal response in BOLD fMRI might be beneficial in the context of rapid event-related designs.

Here, we explore block-design and rapid event-related (rER) BOLD fMRI in awake macaques using visual images of real-world stimuli including human and animal faces, human and animal bodies, objects, and places. In the rER experiment, each stimulus was presented for 0.5 s, and there was a 2.5 s inter-stimulus interval (see Methods). Therefore, we define our event-related design as ‘rapid’ on the basis that the interval between successful stimulus presentations was shorter than the duration of the hemodynamic response function^17^. We selected these stimuli because they have been shown to evoke strong category-selective visual responses in higher-order visual areas in both humans and macaques (see above).

We found clear and strong visual responses and some selectivity to categories, consistent with findings reported in previous studies, in both our block- and rER experiments. However, in the rER experiment, even after censoring scan volumes where our behavioural performance criteria were not met, and after substantial averaging, responses were quite noisy compared to (a) human rapid event-related BOLD fMRI^18^, (b) monkey rapid event-related MION fMRI^14^, and (c) our own block-design BOLD fMRI experiment. We cannot rule out that factors related to the suboptimal performance of our animals may have affected the responses we obtained. Nevertheless, in every volume, our animals maintained eye position, for over half the time, within their fixation window at >82% (see Methods), and eye fixations showed reasonable position stability (see Supplementary Information). To account for our rER fMRI results further, given that collecting additional MRI data using MION was not an option under our study’s project license, we conducted a set of simulations of the BOLD and MION response during rapid event-related experiments. Our results extend previous findings by Leite and Mandeville^19^ by considering all frequencies of stimulus presentation and more conservative assumptions about the MION response shape. Our simulations suggest that the benefits of the greater amplitude of the MION response outweigh the loss of contrast caused by greater temporal smoothing. MION dominated BOLD in functional sensitivity across the entire range of temporal frequencies that matter for event-related experiments and across a wide range of reasonable assumptions about the relative amplitude of MION and BOLD. Together, our experiments using BOLD with block- and rER designs, and our simulations showing improved efficiency of MION than BOLD in rER designs, suggest that future NHP studies aiming to perform BOLD fMRI might benefit from employing a block design, whereas rER NHP studies should consider the use of MION. We conclude that although any two of the three elements (rER, BOLD, NHP) have been shown to work well, the combination of all three is particularly challenging.

## Results

We collected fMRI data while three macaque monkeys (M1-M3) were viewing visual images presented at the centre of a computer monitor.

We first ran a block-design fMRI experiment whose data were also used to define the regions of interest (ROIs) used in our event-related fMRI experiment conducted a few months afterwards. In the event-related experiment, we sought to probe the emergence of selectivity to different object categories in the ventral visual stream, using a stimulus set that has been successfully employed in NHP and human studies previously^14,18,20^. Additionally, given the predominant categorical organisation of face-selective regions on the macaque superior temporal sulcus (STS)^7,21^, we were particularly interested in evaluating the (dis)similarities in the activity patterns^9^ elicited by the individual images in these regions.

### Block-design Experiment

We found strong visual responses for most hemispheres, in the occipital and temporal lobes (figures 1 and 2). Furthermore, we identified anterior, middle, and posterior face-selective regions in the STS (see figure 3 for M1).

**Figure 1:**
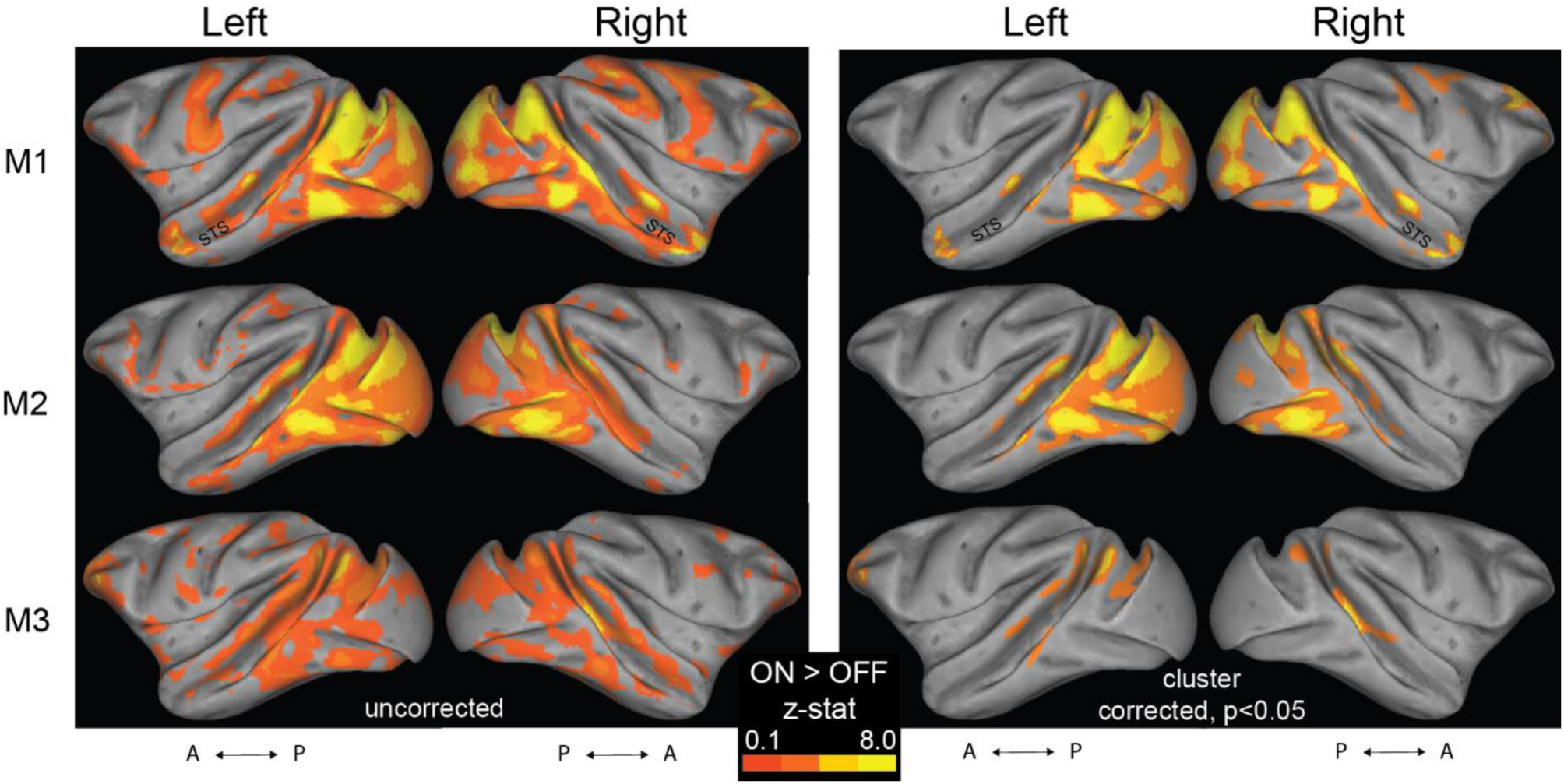
Visual activation (ON>OFF contrast) maps, in the block-design experiment. **Left panel:** Uncorrected data are presented on inflated left and right hemisphere, for the three monkeys (M1-3). For the generation of z-statistic activation maps, see Methods. Results are displayed on surface representations transformed to standard monkey space (MACAQUE-F99; ref^22^). The inflated hemispheres were generated using Caret5 (http://www.nitrc.org/projects/caret/; see ref^23^). A: anterior; P: posterior. **Right panel:** The cluster-corrected z-statistic maps for the three monkeys.

**Figure 2:**
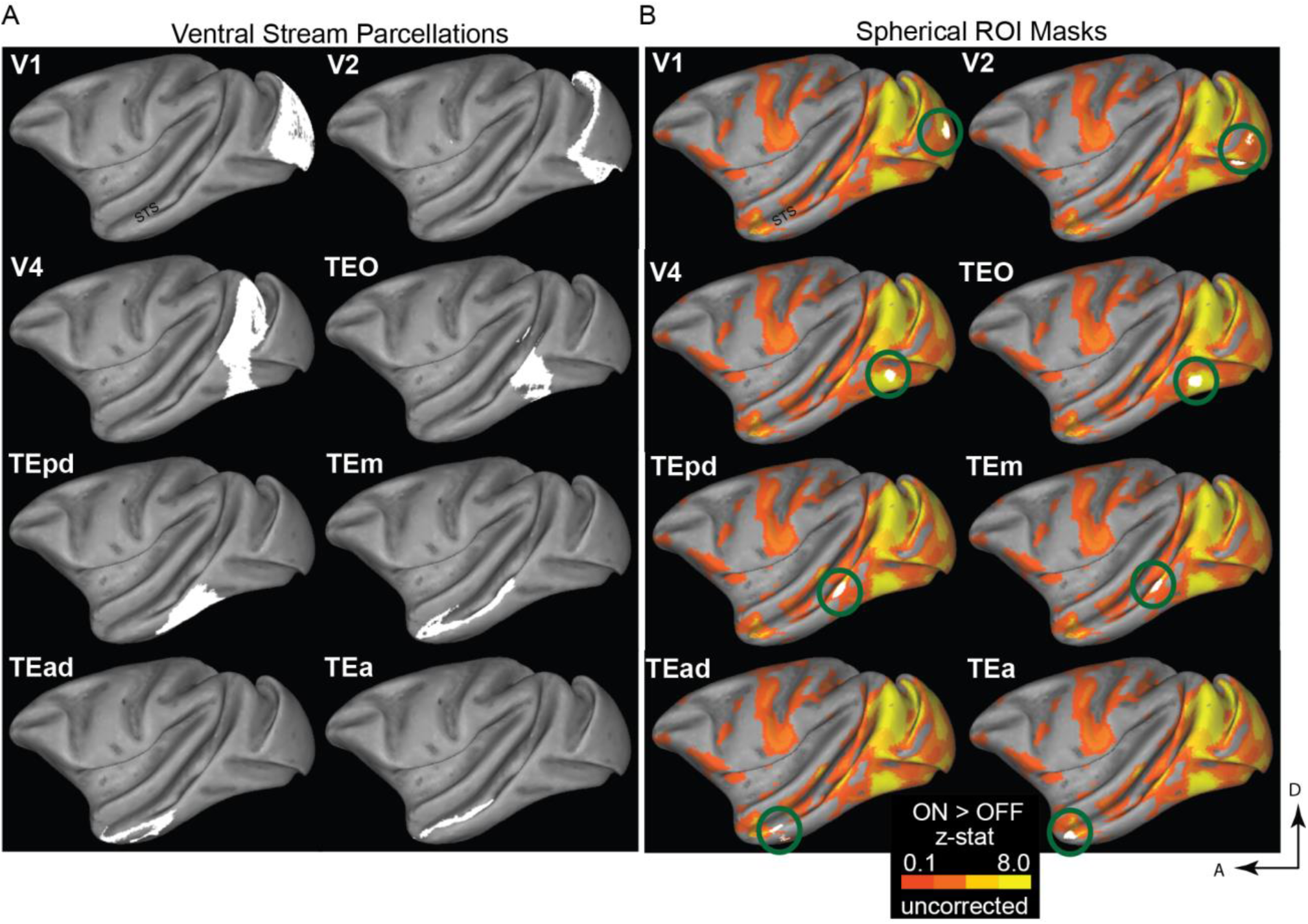
Regions of interest (anatomically-derived) and mask-generation pipeline. **(A)** The regions of the ventral visual stream delineated according to the atlases cortical parcellations described in refs ^24,25^. **(B)** We used the uncorrected ON>OFF results, from the block-design experiment, to generate 2mm radius spherical ROI masks (appearing as white patches, and highlighted with green rings for illustrative purposes) within the original anatomical mask, around the peak ON>OFF voxel, separately for each monkey (see also figure 1). For the ease of representation, the present figure shows the results for M1’s left hemisphere only. D: dorsal; A: anterior.

**Figure 3:**
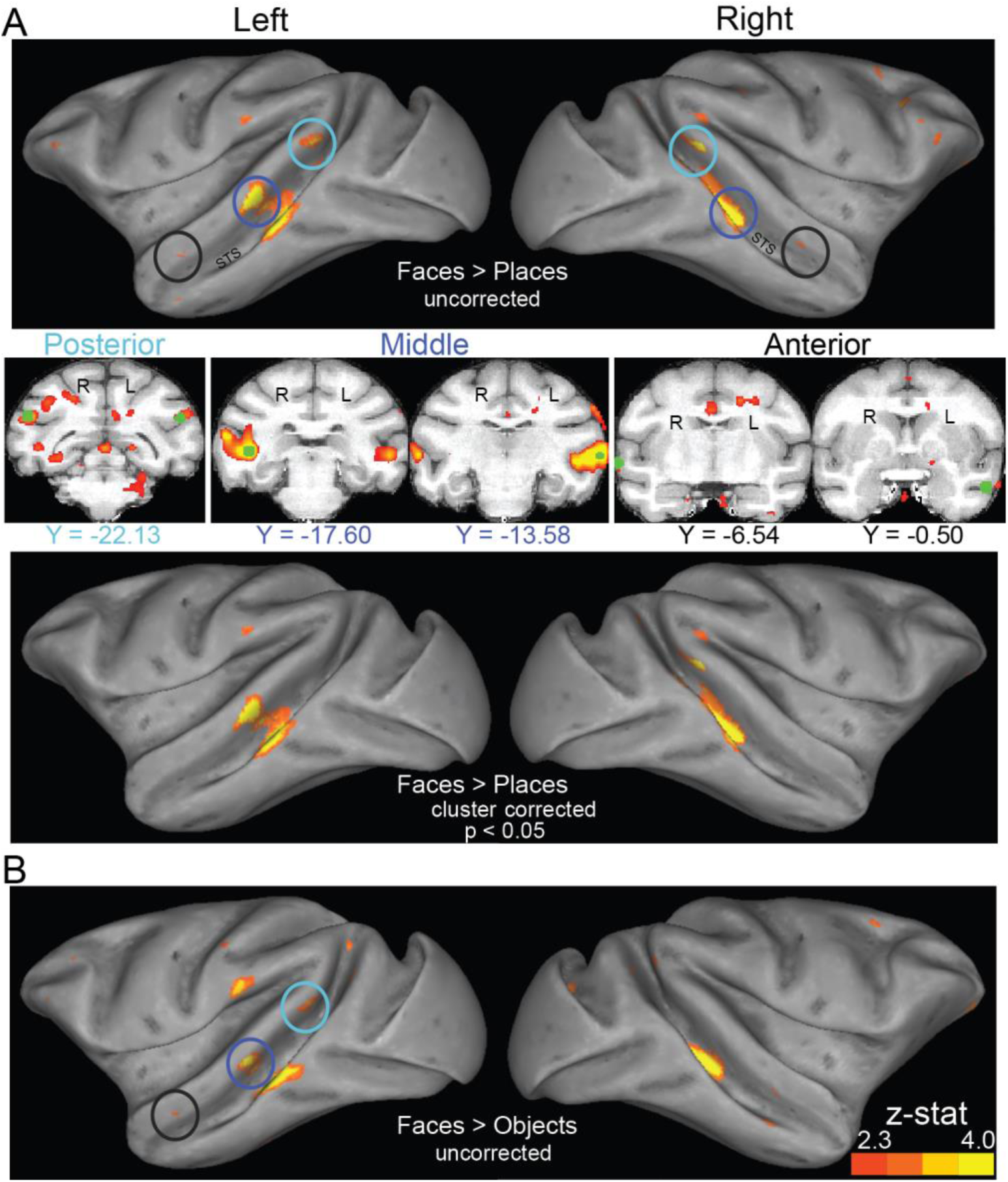
Face-selectivity maps in the block-design experiment. Data are presented on the left and right hemispheres for M1. **A: Activation maps for the faces>places contrast.** Top panel: Uncorrected z-statistic maps displayed on lateral surface representations transformed to a standard monkey brain^22^. In accordance with Tsao et al.^8^, we found regions activated by images of faces in posterior, middle and anterior parts of STS. We generated 2mm radius spheres around the peak face-selective voxels in these posterior, middle and anterior STS (spheres’ rough locations for M1 are highlighted here with cyan, blue and black rings respectively). Middle panel: Coronal-plane views (Y coordinates in MACAQUE-F99 space) of the spherical ROI masks (in green) are also shown for M1. Note that the left hemisphere is shown on the right of each image (radiological display convention). Bottom panel: The cluster-corrected z-statistic maps for M1. By comparison with the uncorrected data, the anterior face patches do not survive the correction for multiple comparisons. **B: Activation maps for the faces>objects contrast:** The STS face-selective regions identified by contrasting faces and objects were either in identical locations or in very close proximity to the regions revealed by the faces>places contrast (especially in the left hemisphere –compare with panel A). Overall, in accordance with ref^4^, the faces>places contrast produced larger regions.

In particular, to localise face-selective regions in each animal, we contrasted face and place stimuli using the data from the present, block-design, experiment. To increase the chances that we captured the face-selective locations in all monkeys, we selected a liberal threshold of z=1.6. Consistent with the ‘face patches’ reported previously^7,8^, we found brain regions with face selectivity in the posterior, middle and anterior parts of STS. As shown in figure 3A (M1’s data; threshold set to z=2.3 for presentation clarity of the face-selectivity found), we identified a large region of face-selectivity that covers the fundus and the lower lip of the middle STS, and likely corresponds to Tsao et al.’s^8^ middle fundus (MF) and middle lateral (ML) temporal face patches. Furthermore, we found a small anterior face region at the fundus of the STS (likely corresponding to the AF in ref^8^) and a small region located more ventrally at the anterior STS (corresponding to the AM in ref^8^). The face-selective voxels we found at posterior STS slightly varied compared to ref^8^: our experiment revealed a posterior face patch in the STS fundus, whereas Tsao et al. reported this to be closer to the STS lip. In the hemispheres where we did not identify a face-selective region (as was the case in M3, where no face-selective voxels were found in anterior or posterior left STS), we used the coordinates from another monkey in our study to generate the mask for the specific missing ROI.

The face-selective regions we found were in close proximity to the ones identified when contrasting faces and objects (figure 3B), further confirming the strong selectivity for face stimuli in macaque STS, as well as providing reassurance that the faces>places contrast we used is appropriate for revealing face selectivity^4^.

### Event-related Experiment: Univariate Analysis

Figure 4 shows percent signal change data across all ROIs for the event-related experiment. The bars depict data averaged across the individual stimuli within each category, then averaged across sessions (all subjects). We found no category-selectivity in early visual cortex V1 or V2. Rather, category-selectivity seems to emerge for the first time at higher levels of visual processing. Specifically, a significant main effect of stimulus category was first observed in area V4 *F(*3,75) = 3.15, *p* =.030). As figure 4A shows, V4 responses to the images of places were greater compared to the responses to the rest of the images. Farther along the ventral cortex, a significant main effect of category (*F(*3,75) = 2.83, *p* =.044) was found in area TEO, where responses to body-part images were greater compared to the other categories. Area TEm showed a significant main effect of category (*F*(3,75) = 2.95, *p* = .041) with a preference to body-parts. We did not find a significant main effect in TEpd (*F*(3,75) = 2.19, *p* = .096), however, responses to body-parts were greater compared to the other categories. Moving more anterior in IT cortex into TEad and TEa, we observed responses lower than baseline to almost all stimulus categories and no significant category-selectivity in either region (*p*’s>0.05).

**Figure 4:**
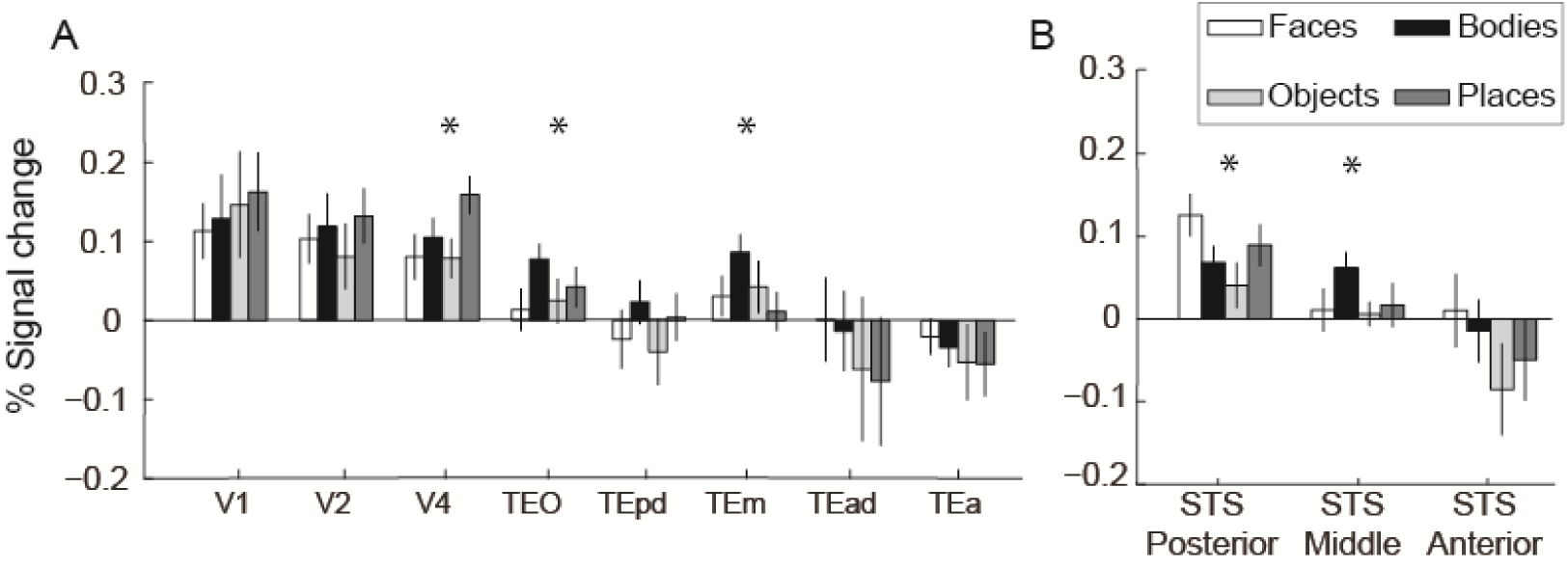
Percent signal change for the event-related experiment. Data are shown across the ventral visual stream (A) and the functionally-defined face-selective regions in STS (B). Results derived from our GLM analysis (contrast: each individual image *versus* baseline). Bars depict data averaged across the individual images within a category, averaged across sessions and across monkeys. The two hemispheres were merged together in a single ROI. Error bars show standard error of the means across sessions.

We also extracted data from the posterior, middle and anterior face-selective ROIs. In the posterior face-selective ROI, we found a significant main effect of category (*F*(3,75) = 4.46, *p* = .006), with greater responses observed to face images. In the middle face-selective STS area, we found a significant main effect of category (*F*(3,75) = 2.95, *p* = .038), with greater responses observed to body-part images. Finally, in the anterior STS area, we found greater responses to faces compared to the other categories, but this did not reach statistical significance (figure 4B).

Finally, similar to the block-design experiment, we contrasted face and place stimuli to generate face-selectivity maps in the event-related experiment. We selected a threshold of 1.6, uncorrected. As shown in figure 5, face-selective regions emerged in reasonable STS locations, however, activations were less strong than in the block-design experiment (figure 3A). Furthermore, contrary to the block-design results, no active voxels survived cluster correction here.

**Figure 5:**
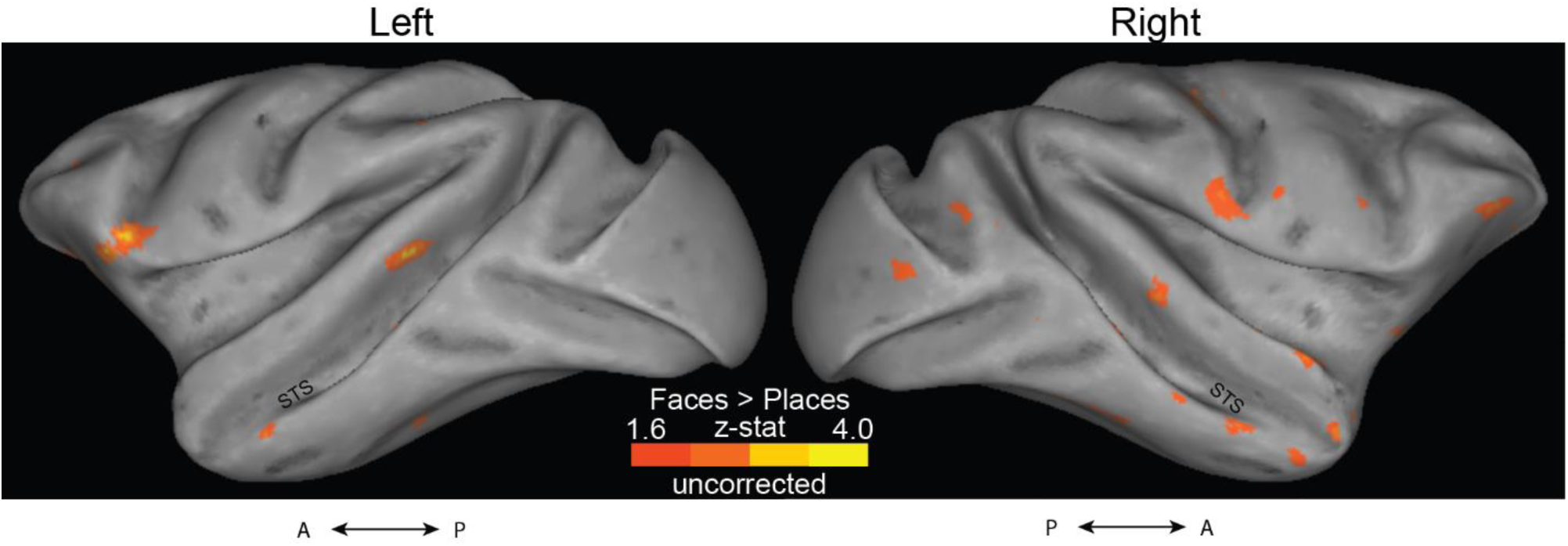
Activation maps for the faces>places contrast, in the event-related experiment. Uncorrected Z-statistic maps for the left and right hemisphere for M1 are displayed on lateral surface representations transformed to a standard monkey brain^22^. Contrary to the block-design experiment (figure 3A), no voxels survived cluster correction here.

### Event-related Experiment: Representational Similarity Analysis

Representational dissimilarity matrices (RDMs)^9^ derived from face-selective regions were visually similar to each other, and RDMs derived from early visual areas were similar to each other, but there were differences across face-selective regions and early visual areas (figure 6). Although there were some dissimilarities in the response patterns between images in the face-selective regions, it is still difficult to come to any conclusions on categorical structure from these data (figure 6 shows an example of an RDM from bilateral V1, posterior face-selective regions, and anterior face-selective regions from M1).

**Figure 6:**
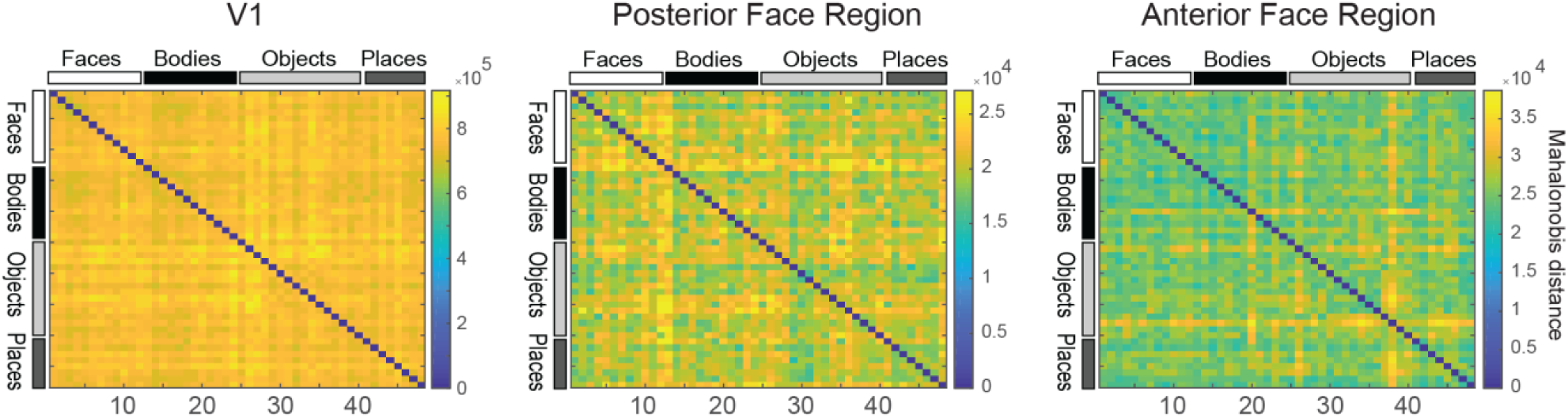
Representational similarity analysis. Examples of representational dissimilarity matrices (RDMs) in bilateral V1 (left), posterior face-selective regions (middle), and anterior face-selective region (right) of M1. RDMs consist of pair-wise cross-validated Mahalonobis distances between all the 48 images we used.

### MION dominates BOLD for simulated event-related monkey fMRI

Our BOLD fMRI rapid event-related response estimates are noisy, suggesting that BOLD rapid event-related designs, although successful in humans, are challenging in monkeys. An important question is whether rapid event-related designs might work better in monkeys when MION is used.

Rapid event-related designs can work with MION^12-14,19^. However, it is unclear how the larger amplitude of the MION response (which helps sensitivity) (see figure 7A) trades off against its larger temporal width (which might reduce the differential sensitivity to fast switching stimuli in rapid event-related designs). Leite and Mandeville^19^ argued on the basis of simulations, that MION more than BOLD benefits from randomization of the stimulus timing, which moves effect energy into lower temporal-frequency bands. Even if high temporal-frequency effects are significantly attenuated in MION fMRI, they could still be stronger than in BOLD fMRI. The power spectrum of the MION model response and linear-model simulations indeed suggests that MION should have greater sensitivity than BOLD in general, i.e. for any type of design^12,19^. However, the MION model used by Leite et al.^12,19^ has a sharp onset, which, on one hand, might not realistically reflect the actual MION response and, on the other, might enable the response model to transmit more high-temporal-frequency information than the actual MION response.

**Figure 7:**
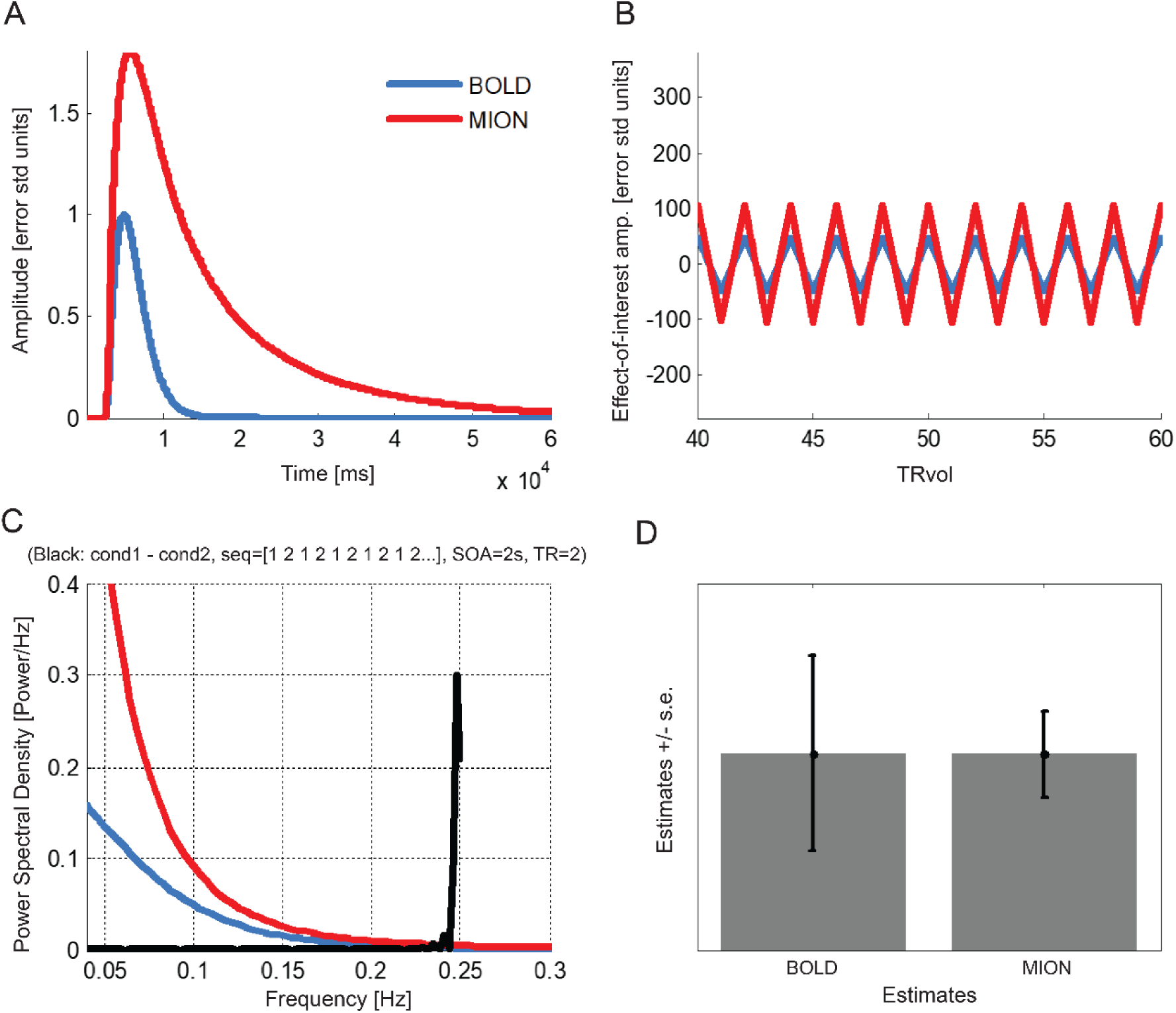
Linear response simulation suggests that MION affords greater sensitivity than BOLD in a fast-switching rapid event-related design. **(A)** Impulse response function models for BOLD (blue) and MION (red). MION has a larger and wider response than BOLD. The BOLD model is from Boynton et al.^26^. The MION model is from Leite et al.^12^, except that the sharp onset has been replaced with the onset of the BOLD model (see figure 8 for the onsets), making the comparison more realistic and conservative with respect to the prospects of MION. Responses are normalized so that the BOLD response peaks at 1. **(B)** Contrast time course. The design consists of two stimuli presented in alternation with a switch every 2s (4s period, 1/4 Hz). The volume TR is 2s. The panel shows the contrast time course, i.e. the stimulus contrast time course convolved with BOLD (blue) and MION (red) impulse response functions. This illustrates that, even for this very fast switching design, where the broad MION response cancels much effect energy, the greater amplitude of the MION response still yields more effect energy than the BOLD response. MION (for equal additive noise) yields greater functional contrast for rapidly switching designs. **(C)** Periodograms of the impulse response functions (red for MION, blue for BOLD) and the stimulus contrast time course (black) show the full-spectrum dominance of MION over BOLD. (**D**) The size of the standard-error bars we expect to obtain with each of the two methods. The greater effect energy for MION translates to substantially smaller standard-error bars.

To assess more conservatively whether MION is theoretically superior to BOLD even for rapid event-related designs, we performed analyses and simulations using a modified version of the Leite et al.^12^ model. We extended Leite et al^12,19^ by replacing the sharp onset of the MION impulse response function with a smooth onset, matching the onset to the BOLD model of Boynton et al.^26^ (figure 7A). We simulated an extremely fast rapid event-related design, in which two experimental conditions (e.g. two stimuli) switch back and forth, with each being presented for the duration of just one acquisition volume (2 s) (figure 7B). Even for this fast-switching design, our simulation shows that MION yields substantially better sensitivity to the contrast between the two conditions (reflected in smaller error bars, figure 7D), essentially replicating the effects predicted by Leite and Mandeville^19^.

To address these questions more generally, we analysed the impulse response of BOLD and MION in the frequency domain (figure 8). We added versions of the BOLD and MION impulse response functions with an even smoother onset than that of Boynton et al.^26^. Results demonstrate that MION has full-spectrum dominance over BOLD.

**Figure 8:**
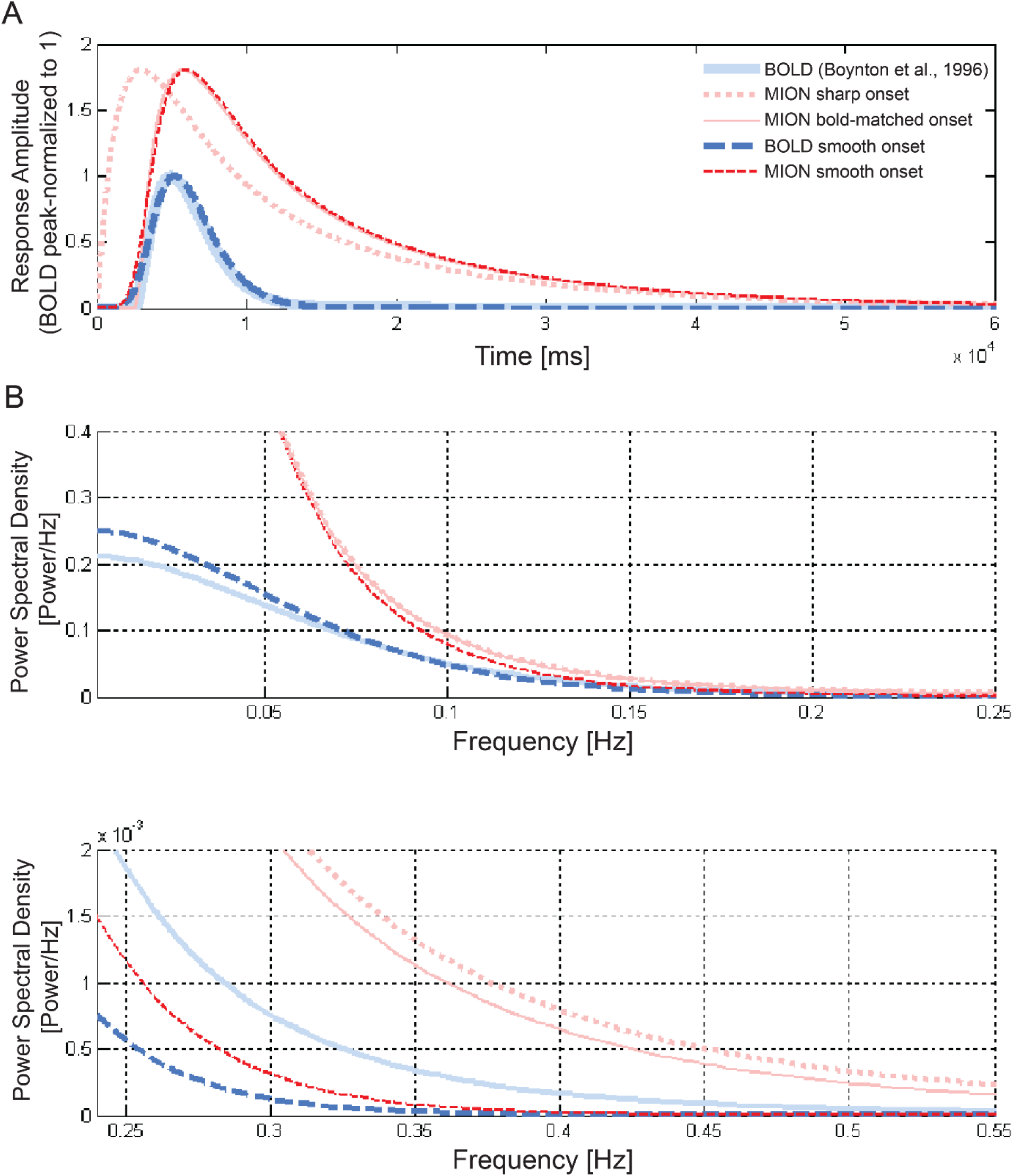
MION affords greater sensitivity than BOLD under conservative assumptions about the onset for arbitrarily rapid event-related designs. **(A)** Different impulse response function models considered for BOLD (blue) and MION (red). The most conservative models (dashed lines in saturated red and blue) have a smooth onset (identical for BOLD and MION), which transmits less high-temporal-frequency effect energy than the Boynton et al.^26^ BOLD model, and much less than the Leite et al.^12^ MION model. **(B)** Periodograms show that for conventional as well as our more conservative smooth-onset models, MION dominates BOLD in terms of its transmission of effect energy across the full spectrum of temporal frequencies. We therefore expect that MION will provide greater sensitivity to effects of interest, no matter how rapid the event-related design.

We tested the robustness of MION’s full-spectrum dominance to changes of the assumed factor by which the peak of the MION response exceeds that of the BOLD response. The previous simulations assumed that the MION response peaks 1.8 times higher than the BOLD response (when the temporal noise is equated). We relaxed this assumption by gradually lowering the peak amplitude of the MION response in the simulation (not shown). MION maintained its full-spectrum dominance over BOLD for rapid event-related designs down to a factor of 1.5 (peak of MION / peak of BOLD) for the smooth-onset variants of both impulse response functions. In sum, the simulations suggest that MION robustly dominates BOLD under conservative assumptions. We expect that MION will yield greater sensitivity no matter what experimental design is used.

## Discussion

We collected BOLD fMRI data from awake behaving macaque monkeys, focusing on the occipital and temporal cortices in two independent experiments: a block-design experiment and a rapid event-related experiment. Each experiment used an independent set of visual images of real-world stimuli. In both experiments, we found visual responses and category selectivity. However, the effects were noisier than expected, especially in the event-related experiment.

In the block-design experiment, we found strong visual responses in the occipital and temporal lobes of all three subjects. Furthermore, we were able to identify bilateral anterior, middle, and posterior face-selective regions for most of the subjects. These face-selective regions were in the regions expected, but less specific than those reported in the literature using MION. For example, Tsao et al.^8^ found six face patches in each hemisphere, where there are two patches for each of the anterior, middle, and posterior parts of the STS. Here, we found correspondence in the anatomical locations, with some subjects showing two patches in each portion of the STS, but these were not easily identifiable in all subjects possibly due to the lower functional contrast of BOLD. However, this could also be because Tsao and colleagues collected more volumes per subject, or had more stimulus repetitions in their block design experiments^8^. We also cannot rule out the possibility that our monkeys’ (who were not fluid-restricted during testing) performance on the task may have negatively affected our observations. For example, on average, we had to exclude some 15% of the collected MRI volumes because monkeys did not sustain viewing within the fixation window, whereas, in the remaining data, fixation eye showed reasonable, yet imperfect, position stability (see Methods and Supplementary Information).

In the event-related experiment, the stimulus-evoked BOLD responses were substantially noisier. Using an ROI approach, we considered ventral stream brain regions from early visual cortices to anterior IT. In early visual areas, we found strong visual responses, but no significant category selectivity. Beyond early visual cortex, we found that category-selectivity begins to emerge. Specifically, we found some category selectivity in V4, TEO, and TEm, as well as in the face-selective regions in the STS. However, as we reach regions in anterior IT such as TEad and TEa, we found no evidence of category-selective responses. This could be related to the relatively weaker fMRI signal found in these regions, but could also be because these regions are more involved in distinguishing identities *within* a particular category (e.g., refs^27-29^ -but see^4^). RSA analyses using noise covariance-normalized distances (crossnobis distances) on face-selective regions found some differences between early visual areas and face regions. The RDMs appeared qualitatively different from each other across areas, but the pattern dissimilarities within a brain region were too noisy for detailed analyses of the representational geometries. Brain activity patterns in early visual areas were strongly dissimilar across stimulus conditions. In the face-selective regions, by contrast, there was only very weak structure suggesting some information about stimulus category. Overall, we found strong visual responses and some category-selectivity in both our block- and event-related designs. However, the data were noisy even after artefact rejection and substantial averaging, which we attribute to the lower contrast-to-noise ratio of BOLD compared to MION, the smaller brains of NHPs, as well as eye-movement- and motion-related artefacts.

Collecting MRI data using MION was not an option under the project license of our study, therefore, we finally performed simulations based on the known response properties of BOLD and MION to compare the response profiles between the two contrast mechanisms. Considering the slower temporal response profile of MION, and previous findings of a more attenuated differential response in MION compared to BOLD at faster rates of stimulus switching^12^, one might expect that BOLD will work better than MION for rapid event-related designs. However, our simulations suggest that at every timescale of stimulus presentation, MION dominates BOLD in terms of sensitivity.

Overall, the stronger BOLD responses we measured in the block-design experiment compared to the rapid event-related experiment suggests that block designs may be a better choice than event-related designs when using BOLD fMRI in NHPs, and our simulations suggest that MION may still be better than BOLD even for rapid event-related designs.

## Methods

### Subjects and Housing

All experimental procedures were performed in accordance with the guidelines and regulations of the UK Animals (Scientific Procedures) Act of 1986. A Project License was reviewed by the University of Oxford Animal Care and Ethical Review Committee and the Home Office (UK) approved and licensed all procedures. Three male macaque rhesus monkeys (M1-M3; mean age: 7 years; mean weight: 12.5 kg) were used in the experiments. M1 and M2 were pair housed and M3 was singly housed, with a 12 hour light/dark cycle (lights on 07:00-19:00). All three animals had unlimited access to water and regular visual contact with human staff. The animals were surgically implanted with an MRI-compatible head post (Rogue Research, Montreal, Canada) in aseptic conditions under general anaesthesia (see ref^30^). After recovery, animals were trained to sit in a primate chair in the ‘sphinx’ position with their heads fixed.

### Training Task and Stimuli

Monkeys were trained to fixate a cue that appeared in the centre of a computer screen. Training took place inside a mock scanner to acclimatize the monkeys to the scanner environment and noise. The images had a size of 11° of visual angle. Monkeys received a smoothie reward for maintaining fixation within a 5° rectangular frame appearing in the centre of each image. The frequency of reward increased over time as the monkey maintained unbroken fixation. Stimulus presentation, eye fixations, and reward delivery were controlled by PrimatePy, a custom-made programme based on Psychopy^31^ (for details on PrimatePy, see^32^). In the scanner, stimuli were presented centrally via an LCD projector onto a rear-projection screen.

In the block-design experiment, we used a subset of the stimuli used by our group previously^33^. Here, our monkeys were presented with images of faces, objects, places, and scrambled versions of objects. In the event-related experiment, the monkeys were presented with a different stimulus set (a subset of the stimuli used in ref^14,18,20^) that consisted of 48 images of human and animal faces and body parts, man-made and natural objects, and places.

### Block-design fMRI procedure

Each image in the ‘ON’ blocks was presented for 0.4 s, with a 0.5 s inter-stimulus interval (ISI). The duration of an ON block was 32 s. ON blocks were interleaved with blank, ‘OFF’, blocks (16 s). Images had a size of 11° of visual angle. A scanning session included a total of 900-2000 volumes, and we collected data in 5 sessions for each monkey. For M1, we collected a total of 7600 volumes; for M2 a total of 6100 volumes; for M3 a total of 7000 volumes.

### Event-related fMRI procedure

On each run (consisting of 117 volumes), each of the images was presented once, in randomized order. Each image was presented for 0.5 s, ISI was 2.5 s, and 30 null trials (blank -isoluminant gray) lasting 2.5 s were interleaved at random time points within a run. A scanning session included a total of 1170-1638 volumes (10-14 runs). For M1, we collected a total of 20358 volumes in 14 sessions; for M2 a total of 12987 volumes in 9 sessions; for M3 a total of 3744 volumes in 3 sessions.

### MR Data Acquisition and Pre-processing

Data for both experiments were collected using a horizontal 3T MRI scanner and a four-channel phased-array receiver coil, together with a radial transmission coil (Wind-miller Kolster Scientific). For the MRI data acquisition, we used an echo planar imaging (EPI) sequence with the following imaging parameters: voxel size=1.5 mm isotropic, repetition time (TR)=2 s, 32 slices, echo time (TE)=29 ms, flip angle=78°.

Raw data were reconstructed offline using a sensitivity encoding (SENSE –see ref^34^) reconstruction method in Matlab, to reduce ghosting artefacts^35^ (Offline SENSE GUI, Windmiller Kolster Scientific, Fresno, CA). To reduce artefacts caused by body motion^36^ we further used motion-correction algorithms^37^ as follows: all volumes within a run were aligned slice-by-slice to the single volume identified as having the least amount of motion (least variance from the mean). The aligned data obtained in the same session were merged into a single 4D NIFTI file, using the Functional MRI of the Brain (FMRIB) Software Library (FSL; www.fmrib.ox.ac.uk/fsl)^38^. In FSL, the 4D data were skull-stripped and subjected to spatial smoothing (full-width half maximum of 3mm), intensity normalisation, and high-pass filtering (cutoff 60 s). Finally, we spatially co-registered the functional data to a standard anatomical template (MACAQUE-F99^22^) using affine transformation.

### Eye-movements and Motion Artefacts

We analysed eye-tracking data to identify and exclude trials where the monkeys broke fixation. Eye-movements were monitored using an MR-compatible LED infrared camera (MRC Systems GmbH, Germany). Eye position was calibrated at the beginning of each session. This calibration procedure was part of the animals’ regular behavioural training in the mock scanner. In the scanner, eye-movements were recorded at 20 Hz, that is, ∼40 samples were obtained in each volume. A volume was excluded if the subject broke fixation for more than half the time in that volume. For each subject, the mean percentage of volumes per session excluded due to broken fixations were: M1=16.3% (standard error of the mean - SEM=2.4%); M2=17.1% (SEM=7.0%); M3=10.0% (SEM=6.7%).

For each session, we identified the volumes containing large motion artefacts (variance two standard deviations greater than the mean of the head motion estimation). The mean percentage of volumes identified per session were: M1=3.0% (SEM=0.3%); M2=4.3% (SEM=0.8%); M3=2.3% (SEM=0.6%). Motion outliers were modelled as nuisance regressors in the main analysis.

### Regions of Interest

The ventral visual stream is considered to be a visual object recognition pathway^39^ accounting for key findings of object-selective responses in monkey inferior temporal (IT) cortex^40,41^. We considered ventral visual stream areas V1, V2, V4, TEO (posterior IT cortex) and TE (anterior IT). We examined subdivisions of TE, that is, posterior TE (TEpd), middle TE (TEm), and anterior TE (TEad and TEa). TEO and TE anatomical masks were delineated according to the macaque cortical parcellations in the Saleem and Logothetis^24^ atlas. V1-4 anatomical masks were delineated according to the cortical parcellations in Van Essen et al.^25^.

Furthermore, recent NHP studies have revealed that the superior and inferior banks of the macaque STS contain several fMRI-identified face-selective regions^4-8^. To reveal such face-selective regions, we contrasted face and place stimuli (from our Block-design experiment data) similarly to previous studies^4,42,43^.

To compare category-selectivity across different parts of the brain in our event-related experiment, we equated the size of all our ROIs^44^. Specifically, for V1, V2, V4, TEO and TE, we created a 2 mm radius spherical mask around the voxel with peak visual activation (ON>OFF contrast in our block-design experiment), within each area (figures 1 and 2). Note that within V4, the spherical masks for the three monkeys were located in the ventral portion of V4^45,46^. For the functionally-defined, face-selective, STS areas, we created a 2 mm radius spherical mask around the peak face-selective voxel (faces>places contrast in the block-design), in the posterior, middle, and anterior STS (figure 3A).

The mask generation pipeline that was applied to all ROIs across both hemispheres for each animal was as follows. Within a given mask, a sphere was generated around the peak visual- or the peak face-selective voxel from our block-design experiment across all sessions within each animal. Before extracting fMRI data from the spherical masks, masks were co-registered to each individual scanning session’s example functional image to align with the functional space of each session. Final spherical ROIs had approximately equal volume (∼30 mm^3^) across animals. We chose spheres of this size so that our spherical ROIs approximately matched the volume of our smallest functionally-defined region (a cluster of face-selective voxels in the anterior STS).

### Event-related fMRI: Data Analysis

We used custom-written code in Matlab to temporally co-register the stimulus presentation times with fMRI volumes and eye-tracking recordings. Statistical analyses were performed using FSL’s FMRI Expert Analysis Tool (FEAT) Version 6.00 by estimating a general linear model (GLM). For each session, each image was modelled as an explanatory variable (EV, i.e., regressor). Monkeys’ head motion outliers (see above) were included in the GLM, as additional EVs of no interest. All EVs were convolved by a hemodynamic response function (HRF) adjusted to reflect the macaque BOLD HRF, which is faster than in humans: we used a gamma HRF with 3 s mean lag and 1.5 s standard deviation (see refs^15,47,48^). We set up one contrast (stimulus > baseline) for each of our 48 images. Z-statistic images arose from EVs according to the following pipeline: each EV in the design matrix resulted in a parameter estimate (PE) image indicating the fit of a waveform model to the data in each voxel. A PE image was converted to a t-statistic image by dividing the PE by its standard error (deriving from the residual noise after the complete model was fit). T-statistic images were then converted to z-statistic images following standard statistical transformations. The beta weight for each stimulus EV, within each ROI, was extracted and converted to % signal change using the *Featquery* tool in FSL. In every ROI, we averaged the data from the two hemispheres.

### Event-related fMRI: Representational Similarity Analysis

To perform representational similarity analysis (RSA)^9^ on face-selective regions and early visual cortex, we extracted the pre-processed fMRI data from these ROIs. ROIs included bilateral anterior, middle, and posterior face-selective regions, and early visual regions bilateral V1 and V2. For the face-selective regions, we used the localization procedure described above, and created new spherical masks with a 5 mm radius. We used larger spherical masks with more voxels for the RSA to improve sensitivity of these analyses, since cross-validated distance measures are based on voxel patterns rather than simply the mean activation in an area. For the early visual regions, we used anatomically-derived masks from ref^25^.

We loaded unsmoothed data from each spherical ROI for each face-selective region in each subject into Matlab and constructed GLMs using custom-written Matlab code. The EVs were modelled as above, with each image modelled as one variable for each run. Each run was modelled separately, in order to perform cross-validation across runs within each session. A GLM-based analysis was performed on each run for each animal and each ROI, which produced a vector of beta weights for each ROI. We used these beta weights to produce representational dissimilarity matrices (RDMs). We used the cross-validated Mahalanobis (or crossnobis) distance, for the distance measure in the RDMs, representing the dissimilarity between two sets of voxel-wise brain patterns^49,50^.

The crossnobis distance was computed as follows:

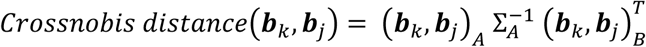

where ***b***_*k*_ and ***b***_*j*_ are vectors of beta weights (fMRI voxel activation patterns) to be compared for image k and j, A denotes the training set and B denotes the test set, Σ_A_ is the noise covariance matrix estimated from the residuals of the GLM for this ROI in the training set A (see ref^50^), and T means transpose. The crossnobis distance was computed between each image in each ROI.

Cross-validation was performed across runs within each session. Given that trials with excessive eye-movements were excluded, not every run included a trial for each image, and therefore it was not possible to use a leave-one-run-out method for computing the crossnobis distance. Instead, we used a split-half approach to estimate the pairwise crossnobis distances. For each session, we randomly assigned half the runs as the training set A and half the runs as the test set B (with one of the runs left out of the analysis when there were odd numbers of runs within a session). This was done 50 times to produce 50 cross-validated distances, and the distances were averaged across cross-validation folds. The same procedure was performed for each session, and the RDMs were averaged across sessions. The noise covariance matrix used was estimated based on the training data. To produce this matrix, we obtained the residuals R estimated from the GLM from an ROI, which is a T (number of time points) x P (number of voxels). The P x P noise covariance matrix can be then estimated by:

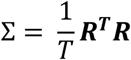

As in the univariate analysis, the RDMs were averaged across hemispheres which gave three RDMs for face-selective regions and three RDMs for early visual areas for each monkey.

## Acknowledgements

This work was supported by a European Research Council Starting Grant ERC-2010-StG 261352 (to N.K.). We would like to thank Dr Urs Schuffelgen for his help and advice on the MRI scans.

## References

1. Downing, P.E., Jiang, Y., Shuman, M. & Kanwisher, N. A cortical area selective for visual processing of the human body. Science 28, 2470–2473 (2001).

2. Epstein, R. & Kanwisher, N. A cortical representation of the local visual environment. Nature 392, 598–601 (1998).

3. Kanwisher, N., McDermott, J. & Chun, M.M. The Fusiform Face Area: a module in human extrastriate cortex specialized for face perception. J. Neurosci. 17, 4302–4311 (1997).

4. Bell, A.H., Hadj-Bouziane, F., Frihauf, J.B., Tootell, R.B.H. & Ungerleider, L.G. Object Representations in the Temporal Cortex of Monkeys and Humans as Revealed by Functional Magnetic Resonance Imaging. J. Neurophysiol. 101, 688–700 (2009).

5. Pinsk, M.A., Desimone, K., Moore, T., Gross, C.G. & Kastner, S. Representations of faces and body parts in macaque temporal cortex: A functional MRI study. Proc. Natl. Acad. Sci. 102, 6996–7001 (2005).

6. Pinsk, M.A. et al. Neural Representations of Faces and Body Parts in Macaque and Human Cortex: A Comparative fMRI Study. J. Neurophysiol. 101, 2581–2600 (2009).

7. Tsao, D.Y., Freiwald, W.A., Knutsen, T.A., Mandeville, J.B. & Tootell, R.B. Faces and objects in macaque cerebral cortex. Nat. Neurosci. 6, 989–995 (2003).

8. Tsao, D.Y., Moeller, S. & Freiwald, W.A. Comparing face patch systems in macaques and humans. Proc. Natl. Acad. Sci. 105, 19514–19519 (2008).

9. Kriegeskorte, N., Mur, M. & Bandettini, P.A. Representational similarity analysis – connecting the branches of systems neuroscience. Front. Syst. Neurosci. 2, 1–28 (2008).

10. Vanduffel, W. et al. Visual motion processing investigated using contrast-agent enhanced fMRI in awake behaving monkeys, Neuron 32, 565–577 (2001).

11. Taubert, J., Van Belle, G., Vanduffel, W., Rossion, B. & Vogels, R. The effect of face inversion for neurons inside and outside fMRI-defined face-selective cortical regions. J. Neurophysiol. 113, 1644–1655 (2015).

12. Leite, F.P. et al. Repeated fMRI using iron oxide contrast agent in awake, behaving macaques at 3 Tesla. Neuroimage 16, 283–294 (2002).

13. Popivanov, I.D., Jastorff, J., Vanduffel, W. & Vogels, R. Stimulus representations in body-selective regions of the macaque cortex assessed with event-related fMRI. Neuroimage 63, 723–741 (2012).

14. Liu, N. et al. Intrinsic structure of visual exemplar and category representations in macaque brain. J. Neurosci. 33, 11346–11360 (2013).

15. Kagan, I., Iyer, A., Lindner, A. & Andersen, R.A. Space representation for eye movements is more contralateral in monkeys than in humans. Proc. Natl. Acad. Sci. 107, 7933–7938 (2010).

16. Kaskan, P.M., Dean, A.M., Nicholas, M.A., Mitz, A.R. & Murray, E.A. Gustatory responses in macaque monkeys revealed with fMRI: Comments on taste, taste preference, and internal state. Neuroimage 184, 932–942 (2019).

17. Buckner, R.L. et al. Functional-anatomic correlates of object priming in humans revealed by rapid presentation event-related fMRI. Neuron 20, 285–296 (1998).

18. Kriegeskorte, N. et al. Matching categorical object representations in inferior temporal cortex of man and monkey. Neuron 60, 1126–1141 (2008).

19. Leite, F.P. & Mandeville, J.B. Characterization of event-related designs using BOLD and IRON fMRI. Neuroimage 29, 901–909 (2006).

20. Kiani, R., Esteky, H., Mirpour, K. & Tanaka, K. Object category structure in response patterns of neuronal population in monkey inferior temporal cortex. J. Neurophysiol. 97, 4296–4309 (2007).

21. Tsao, D.Y. & Livingstone, M.S. Mechanisms of face perception. Ann. Rev. Neurosci. 31, 411–437 (2008).

22. Van Essen, D.C. Windows on the brain. The emerging role of atlases and databases in neuroscience. Curr. Op. Neurobiol. 12, 574–579 (2002).

23. Van Essen, D.C. et al. An integrated software system for surface-based analyses of cerebral cortex. J. Am. Med. Inform. Assoc. 8, 443–459 (2001).

24. Saleem, K.S. & Logothetis, N.K. A combined MRI and histology atlas of the rhesus monkey brain in stereotaxic coordinates. 2nd ed. (San Diego: Elsevier/Academic press. With Horizontal, Coronal and Sagittal series, 2012).

25. Van Essen, D.C., Glasser, M.F., Dierker, D.L. & Harwell, J. Cortical parcellations of the macaque monkey analyzed on surface-based atlases. Cereb. Cortex 22, 2227–2240 (2012).

26. Boynton, G.M., Engel, S.A., Glover, G.H. & Heeger, D.J. Linear systems analysis of functional magnetic resonance imaging in human V1. J. Neurosci. 16, 4207–4221 (1996).

27. Chang, L. & Tsao, D.Y. The code for facial identity in the primate brain. Cell 169, 1013–1028 (2017).

28. Freiwald, W.A. & Tsao, D.Y. Functional compartmentalization and viewpoint generalization within the macaque face processing system. Science 330, 845–851 (2010).

29. Kriegeskorte, N., Formisano, E., Sorger, B. & Goebel, R. Individual faces elicit distinct response patterns in human anterior temporal cortex. Proc. Natl. Acad. Sci. 104, 20600–20605 (2007).

30. Chau, B.K.H. et al. Contrasting roles for orbitofrontal cortex and amygdala in credit assignment and learning in macaques. Neuron 87, 1106–1118 (2015).

31. Peirce, J.W. PsychoPy -psychophysics software in python. J. Neurosci. Methods 162, 8–13 (2007).

32. Joly, O., Baumann, S., Balezeau, F., Thiele, A. & Griffiths, T.D. Merging functional and structural properties of the monkey auditory cortex. Front. Neurosci. 8, 198 (2014).

33. Henriksson, L., Mur, M. & Kriegeskorte, N. Faciotopy –A face-feature map with face-like topology in the human occipital face area. Cortex 72, 156–167 (2015).

34. Pruessmann, K.P., Weiger, M., Scheidegger, M.B. & Boesiger, P. SENSE: sensitivity encoding for fast MRI. Magn. Reson. Med. 42, 952–962 (1999).

35. Kolster, H. et al. Visual field map clusters in macaque extrastriate visual cortex. J. Neurosci. 29, 7031–7039 (2009).

36. Goense, J.B.M., Whittingstall, K. & Logothetis, N.K. Functional magnetic resonance imaging of awake behaving macaques. Methods 50, 178–188 (2010).

37. Kolster, H., Janssens, T., Orban, G.A. & Vanduffel, W. The retinotopic organization of macaque occipitotemporal cortex anterior to V4 and caudoventral to the Middle Temporal (MT) cluster. J. Neurosci. 34, 10168–10191 (2014).

38. Jenkinson, M., Beckmann, C.F., Behrens, T.E., Woolrich, M.W. & Smith, S.M. FSL. Neuroimage 62, 782–790 (2012).

39. Ungerleider, L.G. & Mishkin, M., in Analysis of Visual Behavior (eds Ingle, D.J., Goodale, M.A. & Mansfield, R.J.W.) 549–586 (MIT Press, Cambridge, Massachusetts, 1982).

40. Gross, C.G., Rocha-Miranda, C.E. & Bender, D.B. Visual properties of neurons in inferotemporal cortex of the macaque. J. Neurophysiol. 35, 96–111 (1972).

41. Desimone, R., Albright, T.D., Gross, C.G. & Bruce, C. Stimulus-selective properties of inferior temporal neurons in the macaque. J. Neurosci. 4, 2051–2062 (1984).

42. Rajimehr, R., Young, J.C. & Tootell, R.B.H. An anterior temporal face patch in human cortex, predicted by macaque maps. Proc. Natl. Acad. Sci. 106, 1995–2000 (2009).

43. Rajimehr, R., Bilenko, N.Y., Vanduffel, W. & Tootell, R.B.H. Retinotopy versus face selectivity in macaque visual cortex. J. Cog. Neurosci. 26, 2691–2700 (2014).

44. Hadj-Bouziane, F., Bell, A.H., Knusten, T.A., Ungerleider, L.G. & Tootell, R.B.H. Perception of emotional expressions is independent of face selectivity in monkey inferior temporal cortex. Proc. Natl. Acad. Sci. 105, 5591–5596 (2008).

45. Gattass, R., Sousa, A.P.B. & Gross, C.G. Visuotopic organization and extent of V3 and V4 of the macaque. J. Neurosci. 8, 1831–1845 (1988).

46. Ungerleider, L.G., Galkin, T.W., Desimone, R. & Gattass, R. Cortical connections of area V4 in the macaque. Cereb. Cortex 18, 477–499 (2008).

47. Papageorgiou, G.K. et al. Inverted activity patterns in ventromedial prefrontal cortex during value-guided decision-making in a less-is-more task. Nat. Commun. 8, 1886 doi:10.1038/s41467-017-01833-5. (2017).

48. Fouragnan, E.F. et al. The macaque anterior cingulate cortex translates counterfactual choice value into actual behavioral change. Nat. Neurosci. 22, 797–808 (2019).

49. Diedrichsen, J., Zareamoghaddam, H. & Provost, S. The distribution of crossvalidated Mahalanobis distances. ArXiv arXiv:1607.01371. (2016).

50. Walther, A. et al. Reliability of dissimilarity measures for multi-voxel pattern analysis. Neuroimage 137, 188–200 (2016).

