## Supplementary information for "Rapid event-related, BOLD, NHP: choose two out of three"

**Supplementary Table S1: Eye-tracking data summary statistics within each session, for all monkeys.** Mean and standard deviation of the eye position (degrees of visual angle) across the runs within a session. Although fixations at the centre (zero) of the fixation window (size=5 degrees) were not perfectly sustained throughout, the data suggest that eye position variability within the window was reasonable (<2 deg for the majority of sessions).

| **Scanning Session** | **Horizontal position, Degrees** | | **Vertical position, Degrees** | |
| --- | --- | --- | --- | --- |
| **M1** | Mean | Standard Deviation | Mean | Standard Deviation |
| Session 1 | -0.84 | 1.59 | 0.89 | 1.71 |
| Session 2 | 0.53 | 1.37 | 1.21 | 1.15 |
| Session 3 | -1.05 | 1.46 | 1.07 | 1.51 |
| Session 4 | 0.57 | 1.18 | 0.64 | 1.42 |
| Session 5 | -0.18 | 1.35 | 1.09 | 1.40 |
| Session 6 | 0.57 | 1.35 | 1.78 | 1.13 |
| Session 7 | 0.00 | 1.42 | -0.33 | 1.46 |
| Session 8 | -0.02 | 2.01 | 0.24 | 1.88 |
| Session 9 | -0.67 | 1.46 | 0.11 | 2.04 |
| Session 10 | -0.30 | 1.67 | 0.81 | 1.68 |
| Session 11 | -0.76 | 1.68 | 0.00 | 1.16 |
| Session 12 | -0.64 | 0.85 | 0.23 | 1.05 |
| Session 13 | -0.25 | 1.19 | 0.86 | 1.42 |
| Session 14 | -1.02 | 1.34 | 1.11 | 1.19 |
| **M2** |  |  |  |  |
| Session 1 | -1.00 | 1.08 | 2.02 | 1.31 |
| Session 2 | -1.55 | 1.24 | 0.90 | 1.62 |
| Session 3 | -0.92 | 1.09 | 0.90 | 1.03 |
| Session 4 | -0.40 | 1.06 | -0.69 | 1.55 |
| Session 5 | 0.51 | 1.92 | -0.58 | 2.07 |
| Session 6 | -0.52 | 0.94 | 1.12 | 2.05 |
| Session 7 | -0.36 | 1.52 | 0.46 | 1.99 |
| Session 8 | -0.04 | 1.75 | -0.42 | 1.93 |
| Session 9 | -0.46 | 1.79 | 0.25 | 2.15 |
| **M3** |  |  |  |  |
| Session 1 | 1.13 | 1.40 | 0.86 | 1.27 |
| Session 2 | 0.29 | 1.92 | 1.67 | 1.35 |
| Session 3 | -1.27 | 2.26 | -0.36 | 1.65 |

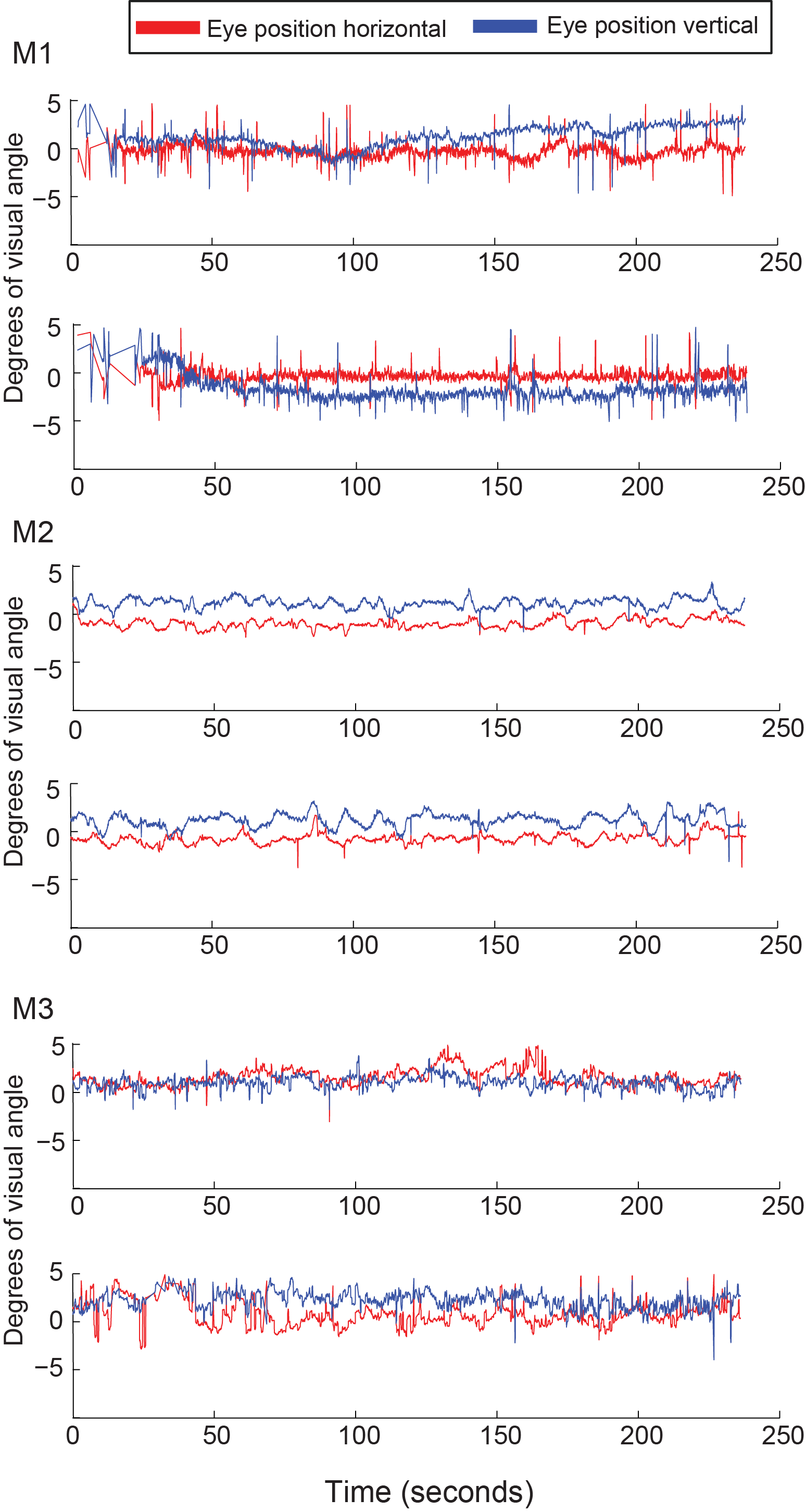

**Supplementary Figure S1:** **Representative subset of the eye-tracking data for all three monkeys.** Each graph shows the eye position on the horizontal and vertical plane, within the animals’ fixating window, expressed in degrees of visual angle, as a function of time. For the purpose of illustrating eye traces in reasonable time chunks, and avoid diffusing the variability between runs due to averaging, each graph shows an individual run. Summary statistics for the entire list of scanning sessions can be found in supplementary table S1.
